# Targeting AKAP13 RhoGEF activity ameliorates pro-fibrotic phenotypes driven by the IPF associated *AKAP13* risk variant

**DOI:** 10.64898/2026.01.21.700846

**Authors:** Bin Liu, James May, Greg Contento, Sara Gangi, Louise Organ, Elizabeth Pyman, Iyobel Kibreab, Lan Zhao, Huayang Yao, Rachel C Chambers, Iain D. Stewart, Anna K. Reed, R Gisli Jenkins, Alison E. John

## Abstract

**Rationale:** Idiopathic pulmonary fibrosis (IPF) is a progressive, incurable scarring disease of the lung. A common genetic variant near AKAP13, a multifunctional scaffold protein that integrates intracellular signalling through its interactions with RhoA and protein kinase A (PKA), has been associated with IPF susceptibility and elevated *AKAP13* mRNA expression in lung tissue from patients. However, its contribution to the pathogenesis of IPF remains unclear.

**Objective:** This study investigates how an AKAP13 variant alters epithelial signalling and evaluates the therapeutic potential of targeting AKAP13.

**Findings:** rs62025270-bearing iHBECs exhibited selective upregulation of AKAP13 isoforms, accompanied by increased cell adhesion and reduced proliferation. Transcriptomic profiling revealed upregulated fibrosis-related genes in rs62025270-bearing iHBECs, including *SAA1*, *FGF2*, *MMP1*, *CTSB*, *COL4A1*, and *CDKN1A*. rs62025270-bearing iHBECs also displayed increased RhoA activation and SMAD2 phosphorylation following LPA stimulation. Furthermore, cells harbouring the AKAP13 variant showed reduced intracellular cAMP levels. Pharmacological inhibition of AKAP13 with A13 reversed the pro-adhesive phenotype and reduced RhoA activation in iHBECs. Moreover, in IPF-derived PCLS, A13 suppressed SERPINE1, CCN2, and MMP7 expression, reduced SMAD2 nuclear translocation, and decreased hydroxyproline levels.

**Conclusions:** Presence of an AKAP13 variant disrupts epithelial homeostasis and promotes pro-fibrotic signalling. Inhibition of AKAP13’s RhoGEF domain with A13 restores epithelial function and attenuates fibrotic activation, supporting AKAP13 as a therapeutic target in IPF.

## Introduction

Idiopathic pulmonary fibrosis (IPF) is a progressive and fatal interstitial lung disease characterised by chronic epithelial injury in a genetically susceptible individual, leading to excessive extracellular matrix (ECM) deposition, and irreversible lung scarring(1-3). IPF has a poor prognosis, with a median survival of only 3–5 years post-diagnosis(4). Current available antifibrotic therapies offer only limited benefit in slowing disease progression, leaving a critical unmet need for more effective treatments.

Recent genome-wide association studies (GWAS) have identified 35 signals associated with increased IPF susceptibility, implicating a wide range of potential molecular pathways; however, the functional consequence of most of these variants remain unknown(5). We have previously reported that the common variant, rs62025270 is associated with increased risk of IPF(6). This variant lies proximal to the gene encoding A-kinase anchoring protein 13 (AKAP13), and associated with increased expression of the *AKAP13* in lung tissue. Consistently, patients with IPF have evidence of increased *AKAP13* mRNA and protein expression in the fibrotic lung(6).

AKAP13 is a multifunctional scaffold protein that integrates cyclic AMP–dependent protein kinase A (PKA) signalling with RhoA activation through its guanine nucleotide exchange factor (RhoGEF) domain(7). AKAP13 (AKAP-Lbc) has been shown to coordinate GPCR-induced signalling in cardiac fibrosis (8-10), where it serves as a RhoGEF and PKA-anchoring scaffold downstream of receptors such as AT1R. AKAP13 suppression markedly attenuates Angiotensin II–induced RhoA activation, myofibroblast differentiation, collagen deposition, and cell migration(8). RhoA is a small GTPase that governs a multitude of cellular processes including actin cytoskeleton dynamics, focal adhesion formation, and cellular contractility(11, 12). In the lung, aberrant RhoA activation promotes fibrosis through avb6 integrin mediated TGFβ activation(13, 14), epithelial cell apoptosis(15), fibroblast activation(16), and mechanical stiffening of the matrix(17). Conversely, cyclic AMP (cAMP) acts as a critical antifibrotic signal, primarily through activation of PKA, which suppresses RhoA activity and attenuates TGFβ-mediated pro-fibrotic signalling(18). A balance between cAMP/PKA and RhoA pathways is, therefore, essential for maintaining alveolar homeostasis and preventing fibrogenesis.

In this study, we investigated the functional consequences of aberrant expression of the AKAP13 rs62025270 variant in IPF pathogenesis. Using CRISPR-Cas9–engineered human bronchial epithelial cells (iHBECs) carrying the rs62025270 variant, alongside isoform overexpression models and pharmacological inhibition of AKAP13’s RhoGEF activity, we examined how variant-induced AKAP13 dysfunction alters epithelial behaviour, fibrotic signalling, and susceptibility to pro-fibrotic stimuli. Findings were further validated in human IPF precision-cut lung slices (PCLS), evaluating the effect of AKAP13 inhibition on fibrotic gene expression and TGFβ pathway activation. Together, our data reveal a novel mechanism by which AKAP13 contributes to lung fibrosis and highlight its potential as a therapeutic target in genetically predisposed individuals.

## Materials and Methods

### Cell culture

Immortalised human bronchial epithelial cells (iHBEC) were cultured in Keratinocyte serum-free medium (KSFM) supplemented with L-glutamine (Gibco), 250 ng/mL puromycin (Sigma), 25 μg/mL Bovine Pituitary Extract (Gibco, #13028014), 0.2 ng/mL human recombinant Epidermal Growth Factor (rEGF) (Gibco), and 25 μg/mL G418 (Life Technologies, Gibco) at 37°C with 5% CO2. For cell starvation, the medium consisted of KSFM with added 250 ng/mL puromycin, and 25 μg/mL G418.

### CRISPR/Cas9 Precision Base Editing

gRNA1 (GGAGACCTGGCTGAGCAGCCGTTTT) and gRNA2 (AGAAATGTCCTTTGAAGGCAGTTTT) targeting the rs62025270 locus for knock-in of the IPF-associated minor allele (A) were designed using Benchling (www.benchling.com). CRISPR/Cas9 constructs were generated with the GeneArt® CRISPR Nuclease Vector OFP Reporter Kit (Life Technologies) according to the manufacturer’s protocol. gRNAs were ligated into GeneArt® vectors, transformed into E. coli One Shot® TOP10, selected on ampicillin agar, and cultured in ampicillin LB broth. Plasmid DNA was purified (QIAprep Spin Miniprep Kit, Qiagen) and gRNA insertion confirmed by Sanger sequencing.

iHBECs were seeded at 2 × 10⁵/well in 24-well and grown to 80% confluency before transfection. For precision base editing, CRISPR nuclease vector containing gRNA were co-transfected with 10 μM single-stranded oligonucleotide (ssOligo) repair template using Lipofectamine™ 3000 (Invitrogen). Cells were incubated with transfection mixtures for 24 h before being sorted using OFP on a FACSAria III (BD Biosciences). Individual iHBECs were seeded into 96-well plates, expanded through successive culture vessels, and genotypes were confirmed using Sanger sequencing.

### Plasmid transfection

Cells were seeded at 2 × 10⁵ cells/well and grown in 6-well plates to 80% confluency prior to transfection. Plasmids encoding FLAG-tagged proto-Lbc, Δ-Lbc and pSR-neo empty vector were delivered using Lipofectamine™ 3000 (Invitrogen) for 48-hour. Transfection efficiency was confirmed at the protein level using immunoblotting as described below.

### Western Blot

Cells were rinsed with ice-cold PBS before being lysed by scraping in buffer containing 100μL/mL Cell Lysis Buffer 10X (Cell Signalling), 100μL/mL cOmplete™ Mini Protease Inhibitor (Sigma-Aldrich), 100μL/mL phosSTOP™ (Sigma-Aldrich), 20μL/mL PMSF (Cambridge Bioscience), 50μL/mL phosphatase inhibitor cocktail (Cambridge Bioscience). Protein concentration was determined using a bicinchoninic acid assay (Fisher Scientific). Cell lysates (30-50 μg) were electrophoresed on Bolt™ 4–12% Bis-Tris gels (Invitrogen) and transferred to polyvinylidene difluoride membranes using the iBlot™ 2 Gel Transfer Device at 25V for 10 minutes (Invitrogen). Membranes were blocked in 5% milk/PBST for 1 h at room temperature, then incubated overnight at 4 °C with primary antibodies: pSMAD2 (0.15ug/ml,138D4, CELL Signalling TECHNOLOGY), SMAD2/3(1:1000, D7G7, CELL Signalling TECHNOLOGY), and AKAP13 (0.375ug/ml, HPA019773, Atlas antibody). After PBST washes, membranes were incubated with HRP-conjugated secondary antibodies (5% milk/PBST) for 1 h and visualized by chemiluminescence (SuperSignal West Femto, Thermo Scientific) using an Odyssey Fc Imager.

### RNA Extraction and cDNA Synthesis

High-quality RNA was extracted from iHBECs using the NucleoSpin™ RNA Mini Kit (Macherey-Nagel, #12373368). Cells (2 × 10⁵/well, 6-well plates) were seeded for RNA extraction. After PBS wash, 350 μL RNA lysis buffer was added, lysates were collected, and RNA was purified with on-column DNase digestion per the manufacturer’s protocol. RNA was eluted in 30 μL RNase-free water and quantified using a NanoDrop spectrophotometer (Thermo Scientific).

For cDNA synthesis, 500 ng of RNA was adjusted to 11 μL with nuclease-free water, mixed with 1 μL oligo(dT)₍₅₀₎ (Promega) and 1 μL 10 mM dNTPs, heated at 65 °C for 5 min, and chilled at 4 °C for 5 min. A master mix containing 1 μL DTT (100 mM), 1 μL SuperScript IV RT (Invitrogen), 1 μL RNase inhibitor, and 4 μL 5× SSIV buffer was added (final volume: 20 μL). Reverse transcription was performed at 55 °C for 10 min, followed by enzyme inactivation at 80 °C for 10 min. cDNA was diluted 1:5 with nuclease-free water and stored at –20 °C.

### nCounter® Fibrosis V2 gene profiling

RNA from iHBECs were used for gene expression profiling using nCounter**®** Fibrosis V2 panel following the manufacturer protocol. Briefly, 50ng of RNA was hybridised at 65 °C with Fibrosis V2 reporter CodeSet and Capture probeSet (BRUKER) for 24 hours before loading into nCounter cartridge and profiled using nCounter SPRINT Profiler (BRUKER). RCC file obtained were uploaded into ROSALIND cloud-based platform for normalisation and downstream analyses.

### xCELLigence Real-Time Cell Analysis (RTCA)

Cell proliferation was monitored using the xCELLigence RTCA system (Agilent) with CIM-Plate 16. 50μL per well pre-warmed KSFM CM+ was added for background measurement. Cells were then seeded at 1 × 10⁵ cells/mL (100 μL/well) and incubated at room temperature for 30 min before placement into the device. Impedance readings were collected every 15 min for 40 h, and cell adhesion and proliferation rates were calculated using RTCA Software 2.0.

### Cell Viability Assay

Cell viability was assessed using the PrestoBlue™ Cell Viability Reagent (Thermo Fisher Scientific) according to the manufacturer’s instructions. Briefly, cells were seeded in 96-well plates at a density of 1 × 10⁴ cells per well in 100 μL of complete growth medium and incubated at 37 °C in a humidified atmosphere containing 5% CO₂ for 24 h to allow cell attachment. The culture medium was then replaced with 100 μL of the treatment solution or complete medium for untreated controls, followed by incubation for 4 h under the same conditions. Subsequently, 12 μL of PrestoBlue reagent was added directly to each well, and the plates were incubated for an additional 24 h at 37 °C. Absorbance was measured at 570 nm with a reference wavelength of 600 nm using a microplate reader.

### Cytotoxicity Detection Assay

Cytotoxicity was evaluated using the lactate dehydrogenase (LDH) assay in accordance with the manufacturer’s instructions. Briefly, cells were seeded in 96-well plates at a density of 1 × 10⁴ cells/well in 100 μL of complete growth medium and incubated overnight at 37 °C in a humidified 5% CO₂ atmosphere. The medium was then replaced with 100 μL of treatment solutions. Cells were treated with 1% DMSO (vehicle control) or 1% Triton X-100 (positive control for maximum LDH release). 100 μL of culture supernatant from each well was transferred to a new 96-well plate, and 100 μL of reaction mixture was added. Plates were incubated at room temperature for 30 min protected from light. Absorbance was measured at 490 nm.

### Intracellular cAMP measurement

Cells were cultured for 48 hours in the presence of adenovirus vectors encoding an Epac1 cAMP sensor, which detects cAMP signalling globally in the cytosol. FRET imaging of epithelial cells was carried out in a FRET buffer solution (144 mM NaCl, 10 mM HEPES, 1 mM MgCl2, 5 mM KCl) that was equilibrated to pH 7.4. Cyan fluorescence protein and yellow fluorescence protein images were acquired using a digital camera attached to the Nikon Eclipse TE2000-U microscope. Cell locations were saved and excited at 430nm wavelength with a 20 images per minute acquisition rate through the Micromanager software. The MultiFRET plugin was used to mark each cell’s ROI and background, enabling individual cell tracking of fluorescence signal changes(19). Baseline signal was recorded for 300 seconds, then 50uM LPA in the fret buffer was added to induce activation of cAMP signalling and recorded for the following 200 seconds. Lastly, 1mL of 10 μM Forskolin and 100 μM 3-isobutyl-1-methylxanthine (IBMX) was added to saturate and measure total cAMP production by fully activating adenyl cyclase and inhibiting phosphodiesterase enzymes.

### RhoA G-LISA Activation Assay

RhoA activity was measured using the RhoA G-LISA Activation Assay Biochem Kit (Cytoskeleton) following the manufacturer protocol. Briefly, iHBECs cultured at 2 × 10⁵ cells per well in 6-well plates were serum-starved for 24 h, then treated with 50 μM lysophosphatidic acid (LPA, Merck) for 2 min. Cells were washed with ice-cold PBS and lysed in 100 μL G-LISA lysis buffer supplemented with protease inhibitors. Lysates were clarified at 10,000 × g for 1 min at 4 °C, protein quantified with Precision Red and diluted to 0.5 mg/mL in lysis buffer. Lysate containing equal amount of protein was mixed 1:1 with binding buffer in duplicate before adding to pre-hydrated assay wells and incubated at 4 °C for 30 min with shaking. Wells were rinsed then incubated with anti-RhoA primary antibody for 45 min, followed by HRP-conjugated secondary antibody for 45 min at 4 °C. Signal was developed with HRP substrate for 10 min at 37 °C, and absorbance was read at 490 nm with plate reader (BioTek).

### TMLC Luciferase Assay

Transformed Mink Lung Cells (gift from Dan Rifkin NYU) were seeded in 96-well plates at 1 × 10⁵ cells/mL (100 μL/well) and incubated at 37 °C, 5% CO₂ for 3 h to allow adherence. Recombinant TGF-β1 (10 μg/mL stock) was diluted to 250, 500, and 1000 pg/mL in growth supplement–free DMEM to generate a standard curve. After adherence, medium was replaced with 100 μL of conditioned medium or TGF-β1 standards in triplicate. Plates were incubated for 16 h, then washed with PBS, and 50 μL of reporter lysis buffer (Promega #E4030) was added. Lysates were transferred to a white opaque plate, and 100 μL luciferase substrate (Promega E1501) was injected using a Luminoskan™ luminometer (Fisher Scientific #15883537) to measure activity. TGFβ activity was calculated from the standard curve relating known TGFβ1 concentrations to luciferase output.

### Human precision cut lung slice (hPCLS) generation

Fresh IPF lung tissue was obtained from the Clinical Research Facility Respiratory Biobank, Royal Brompton Hospital, under ethical approval (NRES 20/SC/0142). To prevent agarose leakage, samples were coated with sodium alginate (3% w/v; Sigma) and cross-linked with calcium chloride (3% w/v; BDH Ltd.) to form a gel seal. Lungs were inflated with sterile 2% low-melting-point agarose (Sigma) in 1× HBSS/HEPES buffer (Life Technologies) at 37 °C and chilled on ice. Tissue cores (8 mm diameter) were prepared using a metal corer (Alabama Specialty Products Inc.) and sectioned at 400 μm thickness using a Compresstome VF-300 microtome. Two hPCLS per well were cultured in 24-well plates with 0.5 mL of RPMI (Gibco) containing 1% penicillin/streptomycin, 0.5% gentamycin, 1 μL/mL amphotericin-B, 25 μg/mL ascorbic acid (Fujifilm), and non-essential amino acids (NEAA, Gibco).

### Immunofluorescence Staining of hPCLS

hPCLS were paraffin-embedded and sectioned at 3 μm thickness. Sections were dewaxed in Histoclear and rehydrated through graded ethanol. Antigen retrieval was performed in citrate buffer (pH 6.0) for 2 min, followed by 40 min cooling to room temperature. Sections were blocked in 3% bovine serum albumin (BSA) in PBS for 1 h at room temperature and incubated overnight at 4 °C with anti–phospho-Smad2 (0.15ug/ml,138D4, Cell Signalling Technology). The following day, sections were washed and incubated with Alexa Fluor 568–conjugated secondary antibody (Invitrogen) for 1 h at room temperature. Nuclei were counterstained with DAPI (1 μg/mL, Roche) for 10 min and slides were washed thoroughly before mounting.

Images were acquired using a Zeiss Axio Observer widefield microscope with a 10× air objective and Zen 2 acquisition software. Nuclear masks were generated using the Cellpose3 algorithm (Python). Phospho-Smad2–positive nuclei were defined as those with a nuclear intensity/(nuclear + 8 μm perimeter) intensity ratio >1, and percentages were calculated accordingly.

### RNA extraction of hPCLS

Total RNA was extracted from human precision-cut lung slices (hPCLS) using TRIzol reagent (Invitrogen) with a Phase-Maker tube (Thermo Fisher Scientific) according to the manufacturer’s protocol. Briefly, 2–3 slices were transferred to a 2 mL Eppendorf tube containing 0.5 mL TRIzol and homogenized using a tissue lyser. Homogenates were transferred to Phase-Maker tubes, adjusted to 1 mL TRIzol, inverted twice, and incubated for 5 min at room temperature. Chloroform (200 μL per 1 mL TRIzol) was added, samples were shaken for 15 s, incubated for 2–3 min, and centrifuged (12,000 × g, 10 min, 4 °C). The aqueous phase was transferred to a fresh tube, and RNA was precipitated with isopropyl alcohol (500 μL per 1 mL TRIzol) and 1 μL glycogen, followed by a 10 min room-temperature incubation. Samples were centrifuged (11,000 × g, 10 min, 4 °C), pellets washed with 75% ethanol (1 mL per 1 mL TRIzol), and centrifuged again (7,500 × g, 5 min, 4 °C). Supernatants were removed, and pellets were air-dried for 10–15 min before resuspension in 30 μL DEPC-treated water.

### Hydroxyproline measurement by high-performance liquid chromatography

Human IPF precision-cut lung slices (PCLS) were cultured for 5 days in the presence or absence of A13. Culture media were collected daily and replaced with fresh medium containing compound. At day 5, supernatants from each slice were pooled per condition for analysis as previously described(20). Briefly, proteins were precipitated by adjusting pooled supernatants to 66% (v/v) ethanol, vortexed briefly, and incubated overnight at 4 °C. Precipitates were collected by filtration through 0.45 μm PVDF filters (MilliporeSigma) and were transferred into glass hydrolysis tubes with 6 M HCl at 110 °C for 16 h to release free amino acids. Following hydrolysis, samples were evaporated to dryness at 100 °C and subsequently derivatized with 7-chloro-4-nitrobenzo-2-oxa-1,3-diazole (NBD-Cl) and quantified by reverse-phase high-performance liquid chromatography (RP-HPLC) using acetonitrile as the organic phase on a C18 (RP-18) column. Hydroxyproline content was used as a quantitative surrogate for secreted collagen.

### Statistical analysis

All quantitative data are presented as median from at least three independent biological replicates. Student’s t-tests were performed for comparisons between two groups, as specified in the figure legends. For multiple conditions or repeated measurements within the same samples, one-sample t-tests or non-parametric Friedman tests with multiple comparisons were applied as appropriate. Statistical analyses were performed using GraphPad Prism. Significance thresholds are indicated in the figures as *p < 0.05 and **p < 0.01.

## Results

### The rs62025270 variant modulates AKAP13 transcripts in human bronchial epithelial cells

The rs62025270 (G>A, 15:85756967) variant was introduced into immortalized human bronchial epithelial cells (iHBECs) using CRISPR–Cas9 and confirmed by Sanger sequencing (Figure 1A). Immunoblotting of protein lysates from variant-bearing cells revealed a prominent protein band of ∼100 kDa in size, considerably smaller than the predicted full-length AKAP13 protein which was expected to be ∼320 kDa (Figure 1B). Region-specific qPCR showed minimal change in the transcript region encoding the including the protein kinase A (PKA)-binding domain compared with wild type (WT) cells, however there was a significant increase in expression of downstream truncated regions, including the Proto-lbc and the RhoGEF domains (Figure 1C). To assess the functional consequences of altered AKAP13 expression, transcriptional profiles of WT and rs62025270-containing cells were analysed using the nCounter® Fibrosis V2 panel (Figure 1D). In comparison with WT cells, variant expressing iHBECs significant upregulated the following genes: serum amyloid A1 (SAA1); fibroblast growth factor 2 (FGF2); matrix metallopeptidase 1 (MMP1); cathepsin B (CTSB); collagen type IV alpha 1 chain (COL4A1), and cyclin-dependent kinase inhibitor 1A (CDKN1A) (Fig 1E-J).

**Figure 1.**
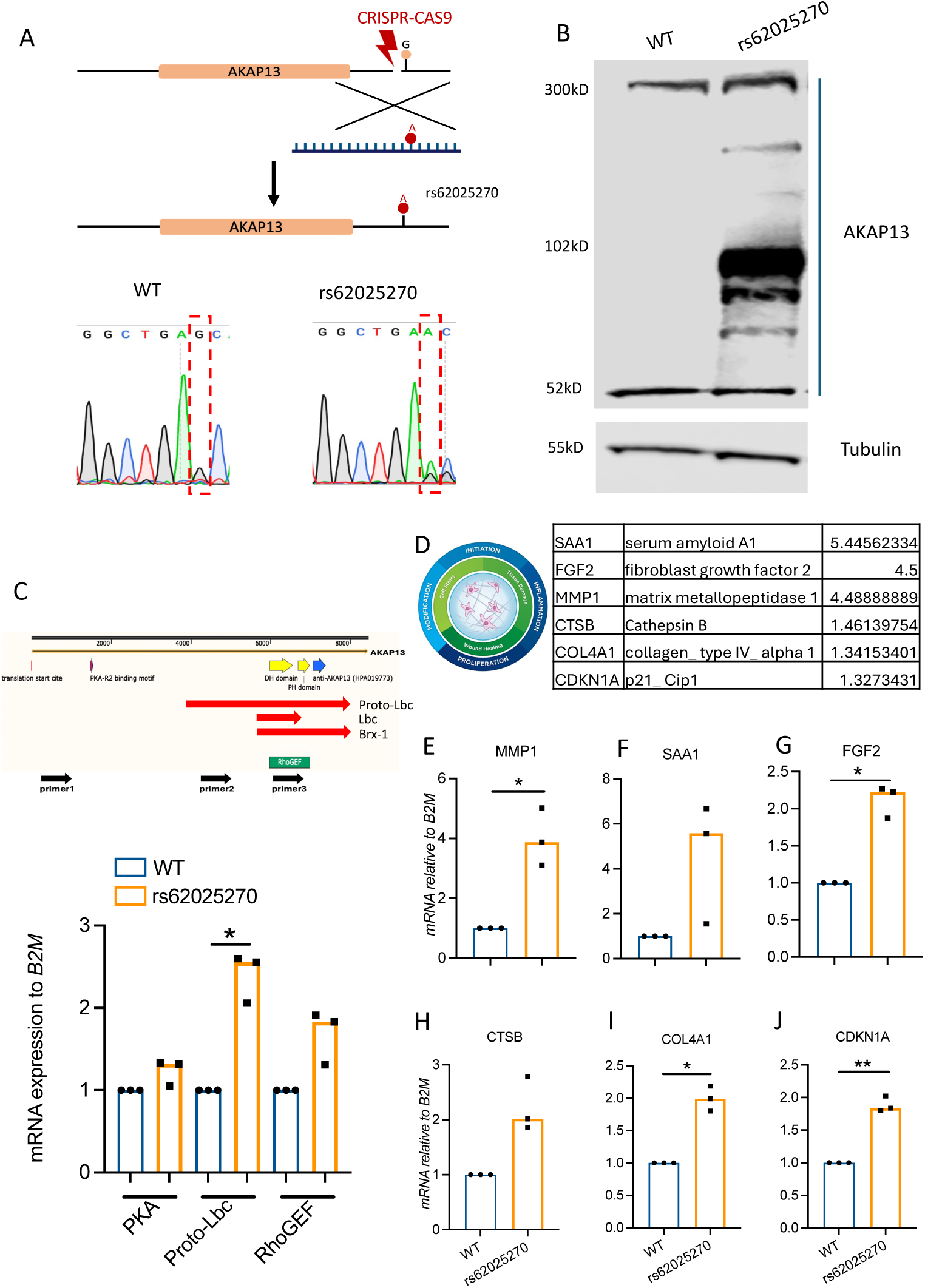
rs62025270 modulates AKAP13 expression in human bronchial epithelial cells. (A) Schematic overview of introducing the rs62025270 (G>A) variant into iHBECs using a CRISPR– Cas9–mediated homology-directed repair strategy. Sanger sequencing confirms the single-nucleotide substitution. (B) Representative immunoblot of AKAP13 and Tubulin protein levels in wild-type (WT) and rs62025270 iHBECs. (C) Region-specific qPCR analysis targeting cDNA regions encoding the PKA (primer 1), proto-Lbc (primer 2), and RhoGEF (primer 3) domains, presented as expression relative to B2M and normalised to WT (n = 3). (D) Top differentially expressed genes (DEGs) identified using the nCounter Fibrosis V2 panel, with adjusted p-value (FDR) < 0.05. (E–J) qPCR validation of selected DEGs: (E) MMP1, (F) SAA1, (G) FGF2, (H) CTSB, (I) COL4A1, and (J) CDKN1A in WT and rs62025270 iHBECs. Data are presented as expression relative to B2M and further normalised to WT (n = 3). Statistical analysis was performed using a one-sample t-test. * p<0.05, ** p<0.01.

### The rs62025270 variant enhances RhoA/TGFβ activation and cellular adhesion

The level of RhoA activation was assessed in wild-type and variant-expressing cells by GLISA assay. Basal RhoA activity was unchanged in rs62025270-bearing cells, however, following stimulation with 50 μM LPA, activity increased by approximately 25% compared to WT (Figure 2A). Real-time impedance monitoring with xCELLigence RTCA revealed that rs62025270 bearing cells had an enhanced cell adhesion rate during the first 5 h (Figure 2B) but reduced proliferation by 40% over 24 h compared with WT cells (Supplemental Figure 1A). Reduced proliferation was confirmed by PrestoBlue assay (Supplementary Figure 1B). Epithelial cell proliferation is inhibited by TGFβ (21) which in turn can be enhanced by LPA-mediated RhoA activation (14). Therefore, the effect of the rs62025270 variant on TGFβ activation was measured. Compared with wild type cells, variant-bearing cells showed augmented SMAD2 phosphorylation following LPA treatment (Figure 2C & D). Furthermore, conditioned media from variant expressing cells showed an increase in TGF-β activity measured by TMLC assay (supplemental Figure 2).

**Figure 2.**
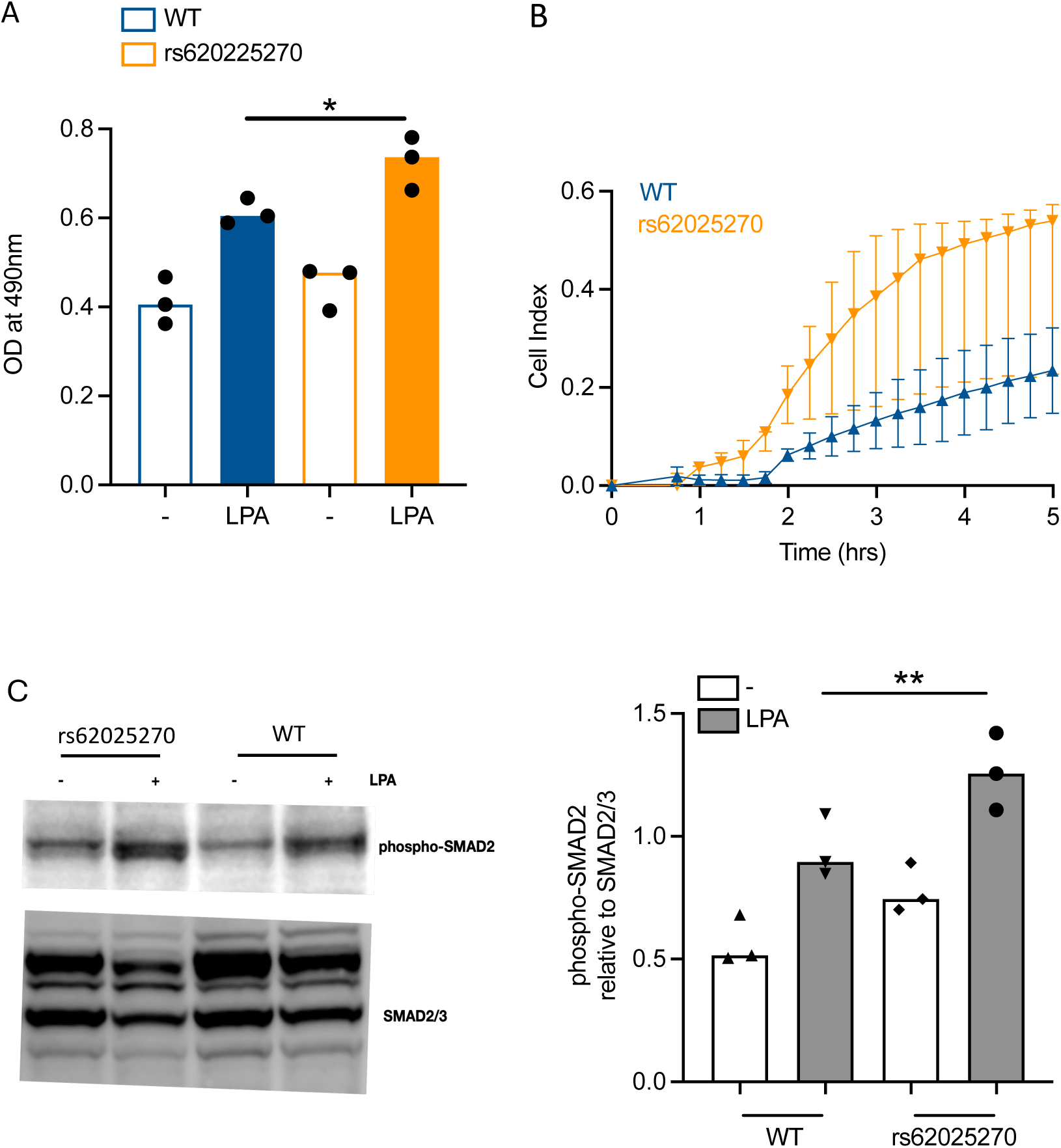
rs62025270 promotes RhoA signalling, leading to increased cell adhesion and TGFβ activation. (A) RhoA activity measured by RhoA G-LISA in wild-type (blue) and rs62025270 (red) iHBECs treated with or without 50 µM LPA for 2 minutes. Data are presented as raw OD values at 490 nm (n = 3). (B) Real-time cell impedance analysis of wild-type (blue) and rs62025270 (red) iHBECs using the xCELLigence RTCA system. Data are shown as cell index over a 5-hour period (n = 3). (C) Representative immunoblot images of phospho-SMAD2 and total SMAD2/3 in iHBECs treated with or without 50 µM LPA for 2 hours. (D) Densitometric quantification of immunoblots shown in (C), presented as phospho-SMAD2 relative to total SMAD2/3 (n = 3). Statistical analysis was performed using a paired t-test between wild type and rs62025270 treated with LPA. * p<0.05, ** p<0.01.

### The rs62025270 variant promotes RhoA activation via increased expression of the proto-lbc

We showed that the rs62025270 variant preferentially induced AKAP13 isoforms lacking the PKA domain (Figure 1C), therefore plasmids encoding the proto-Lbc (an AKAP13 isoform, lacking the PKA domain) or Δ-Lbc (an inactive form lacking RhoGEF domain) (Figure 3A) were selectively overexpressed in iHBECs. Overexpression was confirmed by immunoblotting, with proto-Lbc and Δ-Lbc detected at ∼102 kDa and ∼52 kDa, respectively (Figure 3B). Proto-Lbc overexpression enhanced LPA-induced RhoA activation in iHBECs compared with empty vector, whereas overexpression of Δ-Lbc markedly reduced RhoA activation (Figure 3C). Consistent with the rs62025270 results, overexpression of Proto-Lbc accelerated cell adhesion whereas over-expression of Δ-Lbc diminished cell adhesion compared with the empty vector control (Figure 3D).

**Figure 3.**
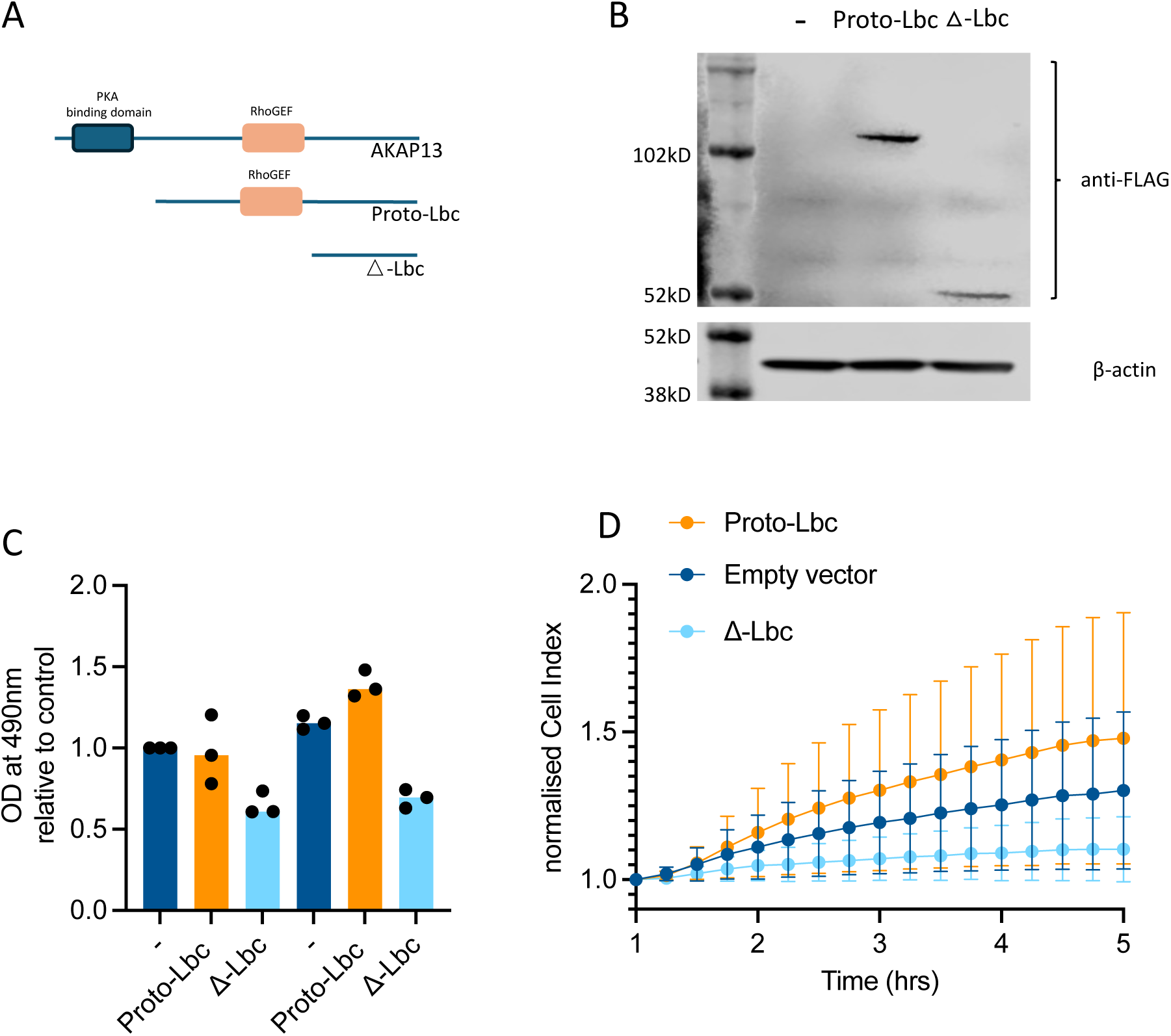
Proto-Lbc overexpression promotes RhoA activation and cell adhesion in human bronchial epithelial cells. (A) Schematic representation of cDNA constructs encoding full-length AKAP13, Proto-Lbc, and △-Lbc. (B) RhoA activity measured by RhoA G-LISA in iHBECs transfected with empty vector, Proto-Lbc, or △-Lbc., followed by treatment with or without 50 µM LPA for 2 min. Data are presented as fold change relative to control (n = 3). (C) Representative immunoblot showing expression of FLAG-tagged Proto-Lbc and Δ-Lbc in iHBECs. (D) Real-time cell impedance measurement of iHBECs using the xCELLigence RTCA system. Data are shown as normalised cell index over a 5-hour period (n = 3).

### The rs62025270 variant reduces cAMP activity

AKAP13 is an A-kinase anchoring protein that is involved in the cAMP signalling, therefore the effect of the rs62025270 variant expressing cells on intracellular cAMP levels was measured using a fluorescence resonance energy transfer (FRET)-based sensor. rs62025270-bearing cells exhibited significantly reduced cAMP levels compared with WT (Figure 4A). To determine whether the effect of cAMP was mediated through the RhoGEF domain of AKAP13 the AKAP13 inhibitor A13 was used. Whilst A13 was unable to restore cAMP to WT levels suggesting that the RhoGEF domain does not contribute to AKAP13 promoting cAMP activity, it was interesting to note a small reduction in cAMP in both wild-type and rs62025270 bearing cells (Figure 4B).

**Figure 4.**
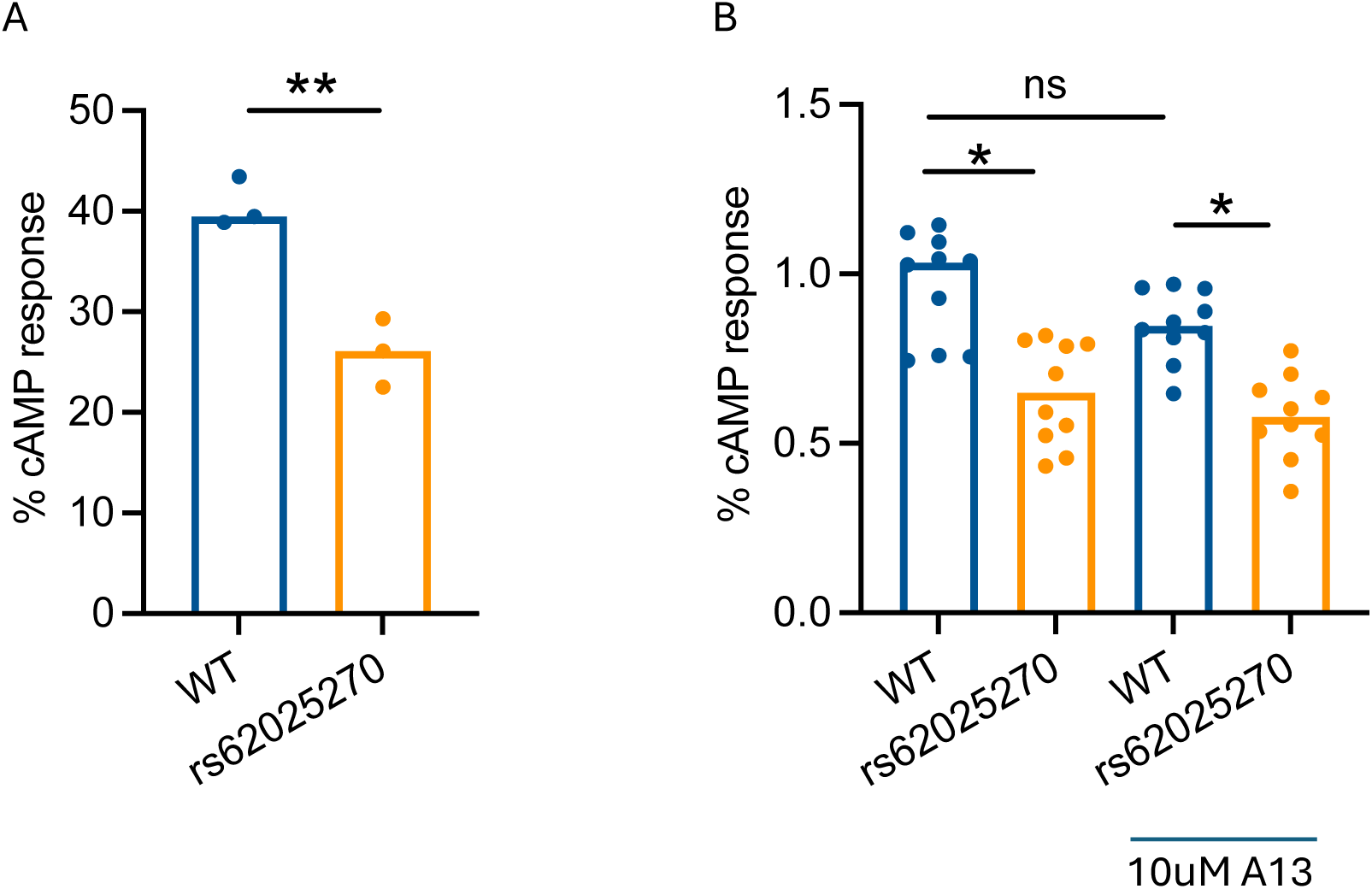
rs62025270 caused intracellular cyclic-AMP (cAMP) reduction, which cannot be restored by A13. (A) Intracellular cAMP levels in iHBECs measured using a FRET-based biosensor. Data are presented as percentage of the FRET signal normalised to the maximum response induced by forskolin (n = 3). Statistical analysis was performed using Student’s t-test. **p < 0.01 (B) Intracellular cAMP levels in iHBECs treated with or without 10 µM A13, as measured by the FRET sensor. Statistical analysis was performed using the Friedman test for multiple comparisons. *p < 0.05

### The AKAP13 inhibitor reduces pro-adhesive phenotype, RhoA and TGFβ activation associated with rs62025270

A13 a small molecule that targets the RhoGEF domain of AKAP13 (22) was used to understand whether targeting this might ameliorate the profibrotic phenotypes observed in rs62025270 bearing cells. A13 inhibited adhesion in a concentration-dependent manner, with 10 μM completely blocking adhesion (Supplementary Figure 3A, B). Notably, 3 μM A13 normalized the elevated adhesion rate of rs62025270 cells to WT levels (Figure 5A). This effect was not due to toxicity as A13 (1–10 μM) did not increase lactate dehydrogenase (LDH) release (Supplementary Figure 3C & D). In contrast, SB431542, which suppresses TGFβ signalling by selectively inhibiting the type 1 receptor ALK5, showed minimal effect on rs62025270 medicated epithelial cell adhesion (supplemental Figure4). Furthermore, 10 μM A13 reduced LPA-induced RhoA activation, restoring rs62025270 associated RhoA activity to WT levels (Figure 5B) and also reduced LPA-induced SMAD2 phosphorylation (Figure 5C & D).

**Figure 5.**
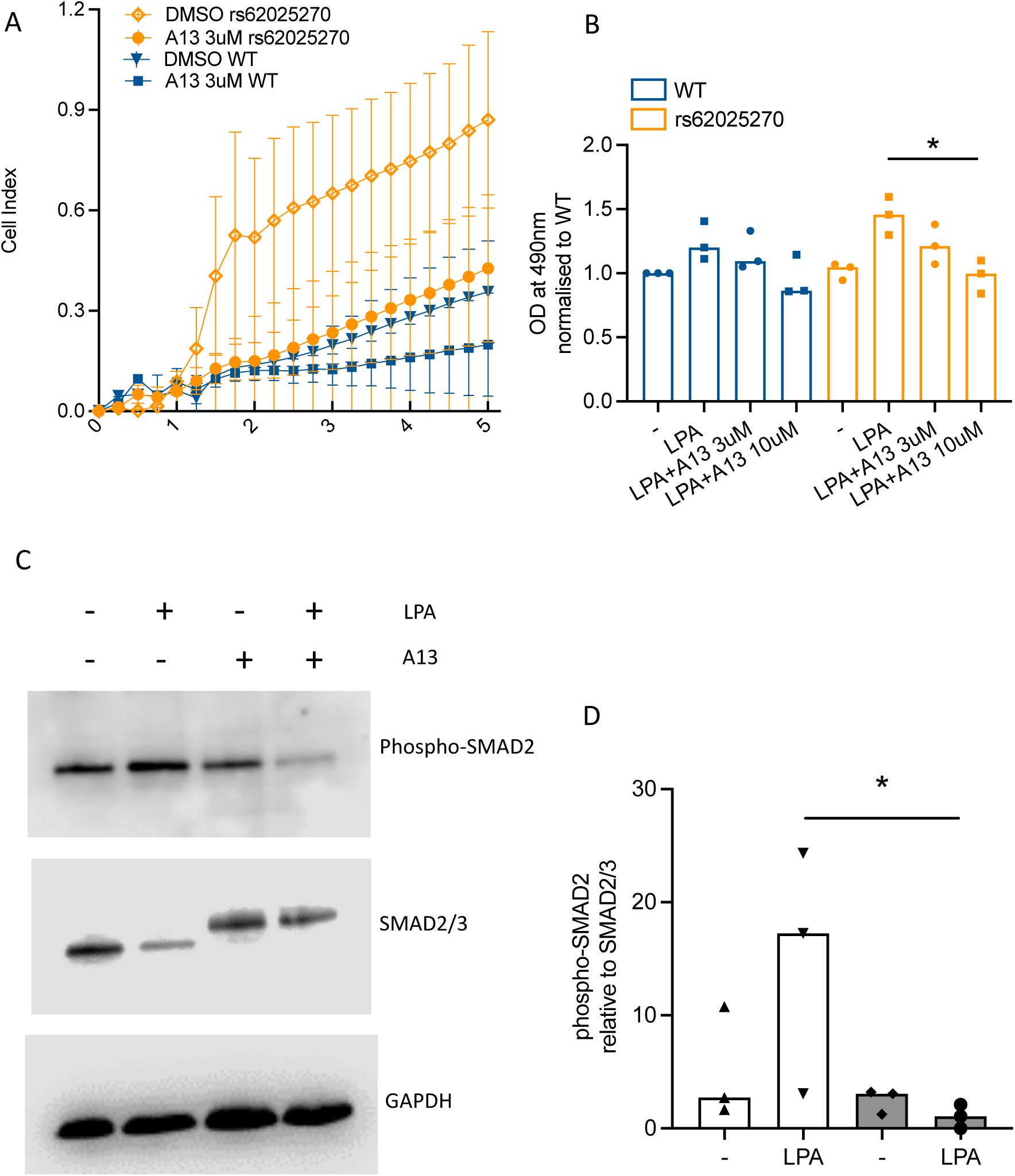
Inhibition of RhoGEF by A13 reverses rs62025270 mediated pro-adhesive phenotype and TGFβ activation. (A) Real-time cell impedance analysis of wild-type (blue) and rs62025270 (red) iHBECs treated with 3 µM A13 or DMSO control using the xCELLigence RTCA system. Data are presented as cell index over a 5-hour period (n = 3). (B) RhoA activity measured by RhoA G-LISA in WT (blue) and rs62025270 (red) iHBECs treated with or without 50 µM LPA for 2 minutes. Data are presented as fold change relative to WT (n = 3). Statistical analysis was performed using the Friedman test for multiple comparisons within genotype. *p < 0.05 (C) Representative immunoblot images of phospho-SMAD2 and total SMAD2/3 in iHBECs treated with or without 50 µM LPA for 2 hours in the presence or absence of 10 µM A13. (D) Densitometric quantification of immunoblots shown in (C). Statistical analysis was performed using the Friedman test for multiple comparisons. *p < 0.05

### An AKAP13 inhibitor has antifibrotic activity in human precision cut lung slices

Finally, we evaluated the antifibrotic potential of A13 in IPF patient–derived precision-cut lung slices (PCLS). 5-day culture of PCLS with A13 significantly reduced the profibrotic gene expression of serpin family E member 1 (*SERPINE1*), cellular communication network factor 2 (*CCN2*), *MMP7*, and fibronectin 1 (*FN1*) in a concentration dependent manner (Figure 6A-D). Consistent with the in vitro findings, A13-exposed PCLS exhibited reduced TGFβ activity, as measured by decreased nuclear phosphorylated SMAD2 (pSMAD2) immunofluorescence staining (Figure 6E & F). 5-day A13 exposure also significantly reduced hydroxyproline content in supernatants at both 3 μM and 10 μM while hydroxyproline levels in lung slice homogenates were similarly affected although to a lesser extent (Figure 6G & H). No toxicity in PCLS was observed with A13 exposure (Figure 6I).

**Figure 6.**
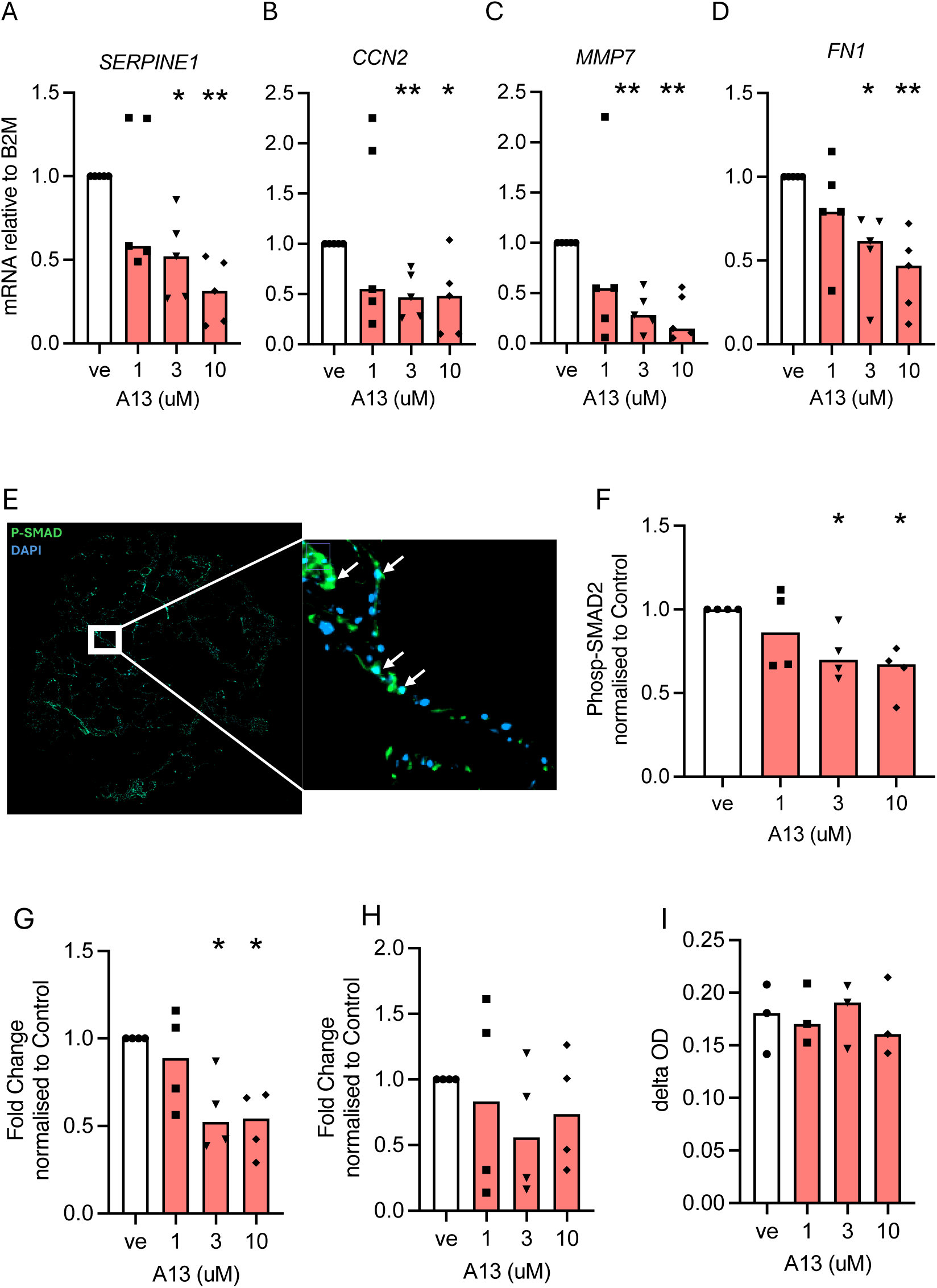
A13 inhibition mitigates fibrosis in IPF precision-cut lung slices. (A–D) mRNA expression of (A) SERPINE1, (B) CCN2, (C) MMP7, and (D) FN1 in PCLS treated with vehicle control or A13 at 1, 3, or 10 µM for 5 days (n = 5). (E) Representative immunofluorescent staining of phospho-SMAD2 (P-SMAD) in IPF PCLS. White arrows indicate cells in which P-SMAD2 (green) colocalises with DAPI (blue). (F) Percentage of nuclear P-SMAD–positive cells normalised to vehicle control (n = 5). (G–H) Hydroxyproline content measured by HPLC in (G) supernatant and (H) tissue slices. Data are presented as arbitrary units normalised to slice weight (n = 4). (I) LDH levels in supernatant from PCLS treated with or without A13 (n = 3). *p < 0.05, ** p < 0.01, one-sample t-test.

## Discussion

In this study, we describe the functional consequences of the rs62025270 variant that has been associated with elevated *AKAP13* expression and is a causal variant for IPF. Using a CRSIPR-Cas9 based approach, introduction of the rs62025270 variant into human bronchial epithelial cells led to enhanced RhoA and TGFβ activation, promoted epithelial adhesion, reduced epithelial proliferation, and lowered intracellular cAMP activity. We show *in vitro* that this variant selectively enhances expression of AKAP13 isoforms lacking the PKA domain but retaining the RhoGEF domain. This altered profile shifts the intracellular signalling balance toward profibrotic RhoA activation and away from anti-fibrotic cAMP activity. Furthermore, we show that specific inhibition of the RhoGEF domain of AKAP13 can suppress profibrotic makers and hydroxyproline production in PCLS from patients with IPF.

Our transcriptomic profiling revealed that the rs62025270 variant elicits a broader shift toward a pro-fibrotic transcriptional state. Specifically, induction of *SAA1*, an acute-phase reactant(23) that has previously been associated with IPF disease severity(24), may suggest variant-bearing cells are undergoing cellular stress. Transcriptional upregulation of the proteolytic enzymes, *MMP1* and *CTSB*, were also observed in the variant-bearing cells, indicating hyperactive extracellular-matrix turnover, which aligns with profibrotic phenotypes observed in pulmonary fibrosis(25, 26). In addition, upregulation of *COL4A1*, a basement-membrane component, may reflect attempts at matrix restoration or remodelling, although the functional consequences of COL4A1 induction in epithelial cells remain unclear and warrant further investigation. Increased *CDKN1A* together with the proliferation suppression supports a shift toward cell-cycle arrest and senescence in variant-bearing cells, a phenotype documented in IPF pathogenesis (27, 28). The induction of *FGF2,* a growth factor important for lung development(29), further suggests activation of epithelial repair. While FGF2 can support acute epithelial recovery, its chronic activation may alter epithelial–mesenchymal signalling, moreover the fact that Nintedanib inhibits FGFR kinases(30) raises the possibility that these cells may respond favourably to existing antifibrotic therapy. Functionally, rs62025270 variant-expressing cells did not exhibit higher basal RhoA activity, but RhoA was increased following LPA stimulation, illustrating the importance of environmental “second hits” in promoting fibrosis in people with common variants(31). Variant-expression cells also showed accelerated cell adhesion, a phenotype reproduced by overexpression of proto-Lbc (the AKAP13 isoform lacking the PKA domain) and suppressed by expression of Δ-Lbc (the RhoGEF-deficient mutant). These findings indicate that the RhoGEF activity of PKA-deficient AKAP13 isoforms is a critical driver of the rs62025270 variant phenotype.

Importantly, the rs62025270 variant not only altered LPA-induced RhoA activity but also amplified LPA-induced TGFβ pathway activation, as shown by increased Smad2 phosphorylation and elevated TGFβ bioactivity in conditioned media. This mechanistic link is consistent with prior evidence that RhoA–ROCK signalling enhances TGFβ activation by modulating cytoskeletal tension and integrin-mediated release of latent TGFβ complexes (14, 32). In addition, AKAP13 functions as a scafford protein for cAMP-dependent protein kinase (PKA) activation. Previous studies have shown that AKAP13 anchored PKA inhibits AKAP13 RhoGEF activity by phosphorylating AKAP13 at the RhoGEF binding site (33). Conversely, recruitment of phosphodiesterase4 (PDE4) to AKAP13 promotes cAMP degradation, resulting in reduced PKA activation(33). Consistent with these observations, we observed reduction in intracellular cAMP in rs62025270 cells, suggesting that loss of PKA-mediated counter-regulation may contribute toward pro-fibrotic signalling. This is particularly relevant given that impaired cAMP signalling has been implicated in IPF pathogenesis, where cAMP modulates fibrosis by affecting fibroblast activation, extracellular matrix production(10, 34).

Pharmacological inhibition of AKAP13 RhoGEF activity using A13 reversed several rs62025270 variant-induced phenotypes. A13 at 3uM effectively normalised rs620250270 variant mediating cell adhesion rates and suppressed LPA-induced RhoA activation to the level of wild type iHBECs. Interestingly, direct inhibition of TGFβ signalling with ALK5 inhibitor showed minimal effects on cell adhesion, suggesting additional mechanisms medicated by RhoA activation that operate independently of TGFβ. Furthermore, 5-day A13 treatment *ex vivo* reduced pro-fibrotic gene expression and *de novo* collagen production in precision-cut lung slices (PCLS) from IPF patients, suggesting that inhibiting the RhoGEF activity of AKAP13 might be useful therapeutic strategy to treat fibrotic lung diseases. Given the impact of the rs62025270 on cAMP signalling it may be possible that targeting the RhoGEF activity of AKAP13 could provide enhanced benefits when combined with currently available anti-fibrotic drugs that modulate this pathway, such as Nerandomilast (35) and Treprostinil (36). Indeed, the finding that the rs62025270 may impair intracellular cAMP activity raises the prospect of stratifying responses to Nerandomilast and/or Treprostinil therapy by *AKAP13* genotype. We would hypothesise that patients with the presence of the rs62025270 would exhibit an enhanced therapeutic response compared with those patients without the variant.

A limitation of this study is that our mechanistic analyses were performed in bronchial epithelial cells, which, although relevant for modelling airway injury responses, may not fully recapitulate the biology of alveolar epithelial cell in IPF. Future work should extend these findings to alveolar type II cells, where epithelial–mesenchymal crosstalk may play a central role in fibrosis progression. Additionally, in vivo studies will be required to evaluate the safety, pharmacokinetics, and efficacy of AKAP13 RhoGEF inhibition in fibrotic lung disease.

In summary, we demonstrate that the IPF-associated rs62025270 variant promotes a shift in AKAP13 isoform expression that favours pro-fibrotic RhoGEF-dependent RhoA activation and impairs anti-fibrotic cAMP activation. Pharmacological inhibition of this pathway attenuates fibrosis-related gene expression, matrix deposition, and TGFβ signalling in human lung tissue. These findings provide functional genetic insight into the rs62025270 risk allele in IPF and establish AKAP13 RhoGEF activity as a promising therapeutic target in genetically susceptible individuals.

## Acknowledgement

This study was funded by a Medical Research Council Programme Grant (MR/V00235X/1) to R.G.J. RGJ was funded by an NIHR Research Professorship (RP-2017-08-ST2-014). B.L. is a research fellow funded by Action for pulmonary fibrosis. I.D.S. is an advanced research fellow funded by Rayne Foundation. Infrastructure support for this research was provided by the NIHR Imperial Biomedical Research Centre (BRC). The authors acknowledge Clinical Research Facility (CRF) Respiratory Biobank, Royal Brompton Hospital for their collection of tissue and facilitation of sample transfer.

## Author contributions

B.L., J.M., S.G., L.O., E.P., contributed to in vitro experiments, tissue sample collection, PCLS generation, and data analysis. G.C. performed hydroxyproline measurement. B.L. and I.K. performed intracellular cAMP measurement using FRET sensor. L.Z., R.C.C., critically reviewed and revised the manuscript. B.L., A.E.J., and R.G.J. conceptualized the study and drafted the manuscript. A.E.J. and R.G.J. supervised the project, coordinated collaborations, and finalized the manuscript. All authors reviewed and approved the final version of the manuscript.

---

**Supplemental Figure1.**
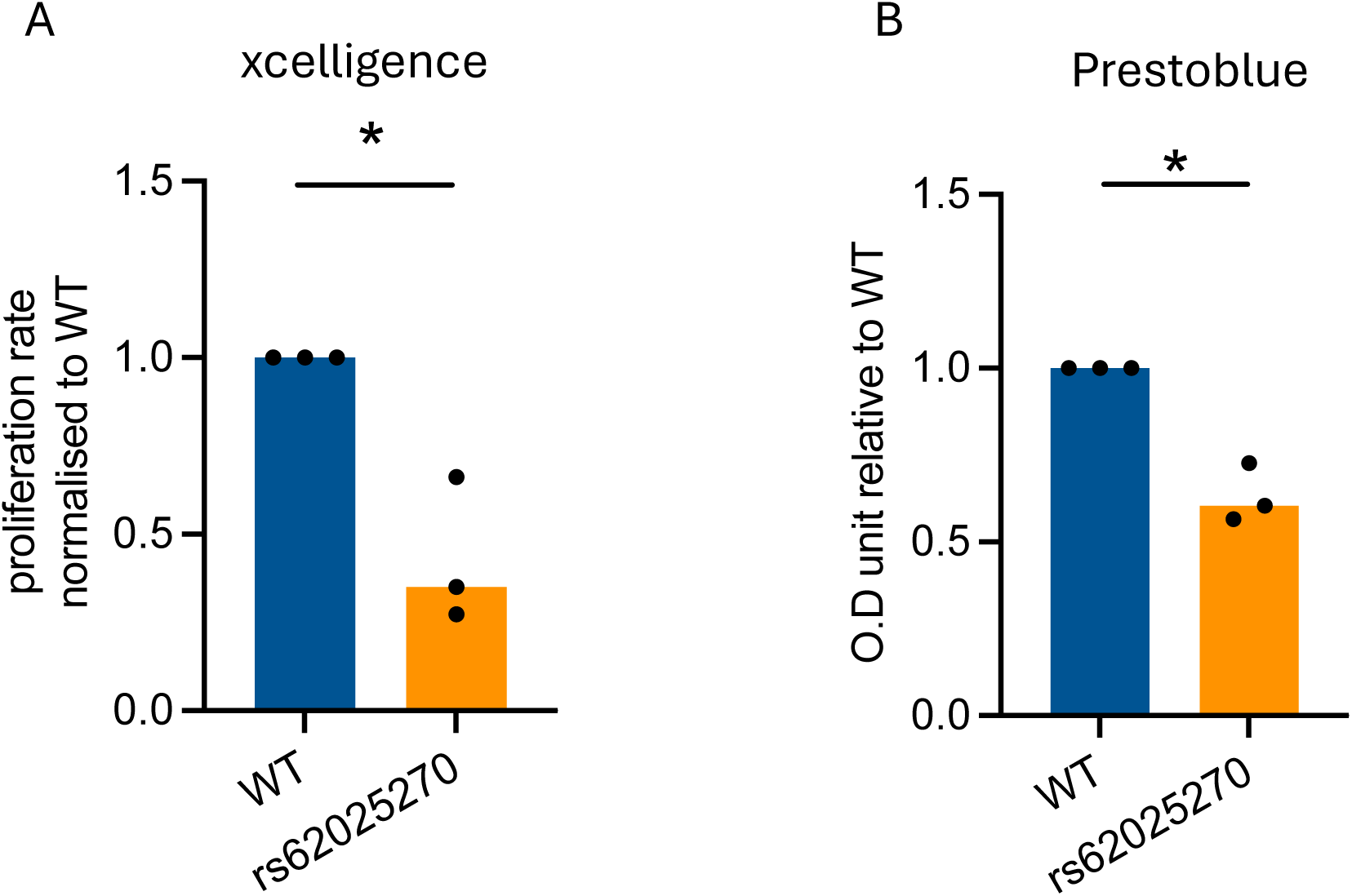
cell proliferation measured by xcelligence and Prestoblue. (A) Cell proliferation measured by xcelligence RTCA and normalised to wild type over 24 hours. (B) Cell proliferation measured by Prestoblue over 24 hours and presented as O.D value. Statistical analysis was performed using a paired t-test between wild type and rs62025270 treated with LPA. * p<0.05

**Supplemental Figure2.**
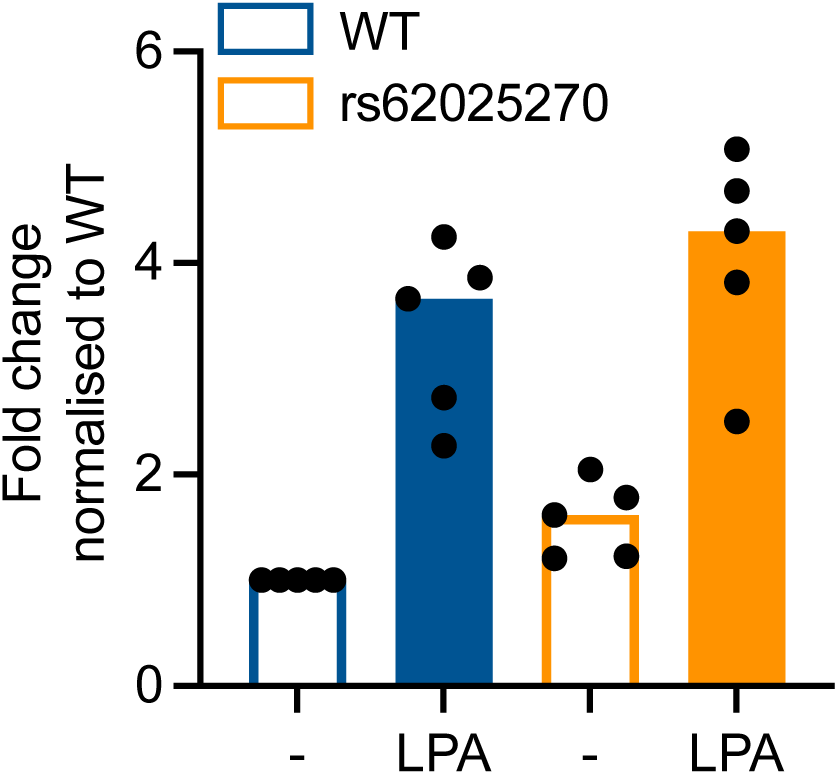
TGFβ activity measured by TMLC reporter assay. TGFβ activity in conditioned media measured using a TMLC reporter assay. Data are presented as fold change relative to vehicle control and further normalised to WT (n = 5).

**Supplemental Figure3.**
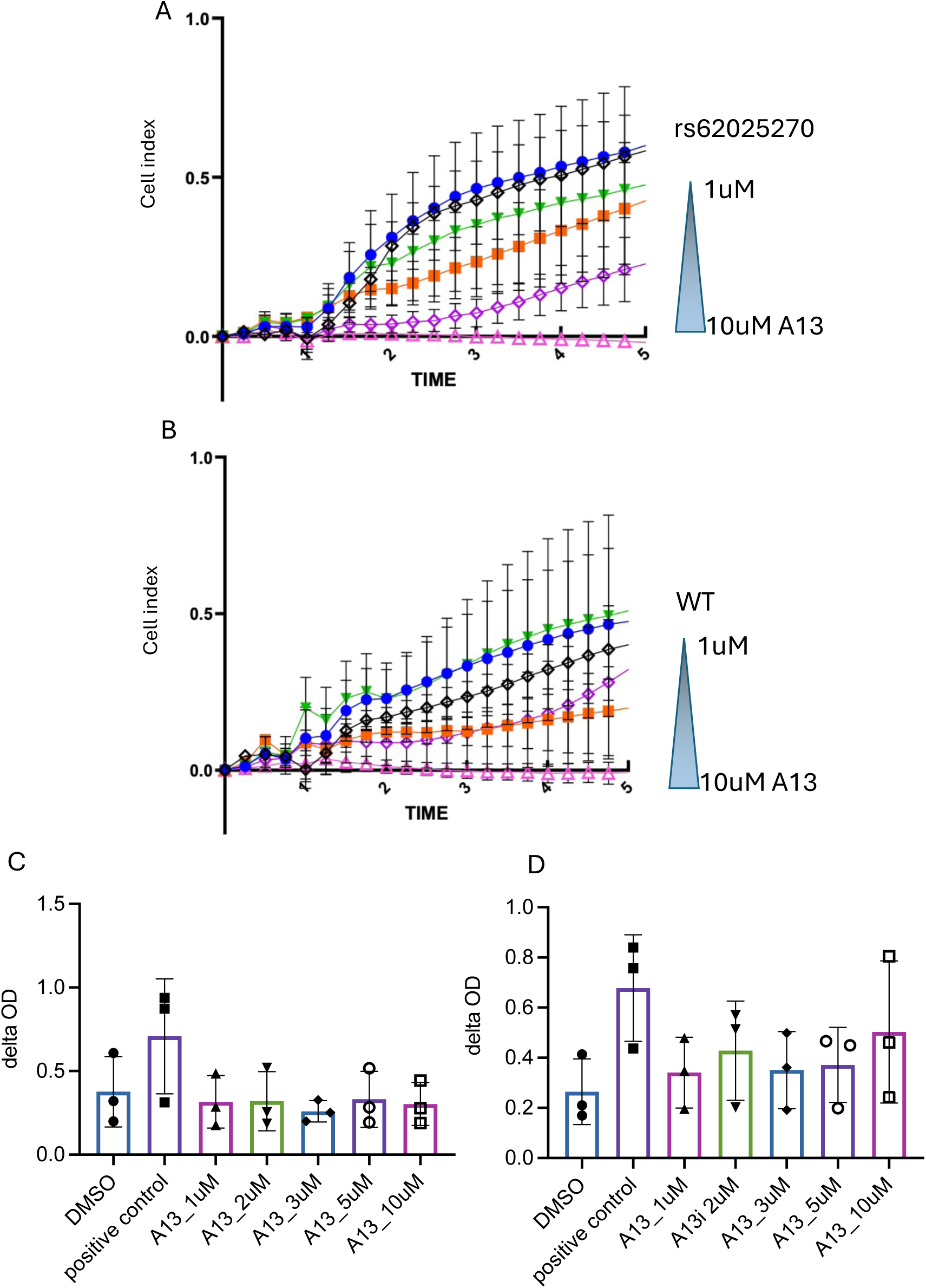
Real-time cell impedance and cytotoxicity assessment of A13-treated iHBECs. (A–B) Real-time cell impedance measurements of (A) rs62025270-bearing and (B) WT iHBECs treated with A13 (1–10 µM) or DMSO control using the xCELLigence RTCA system. Data are presented as cell index over a 5-hour period (n = 3). (C–D) Lactate dehydrogenase (LDH) levels in (C) rs62025270-bearing and (D) WT iHBECs treated with A13 (1–10 µM) or DMSO control (n = 3).

**Supplemental Figure4.**
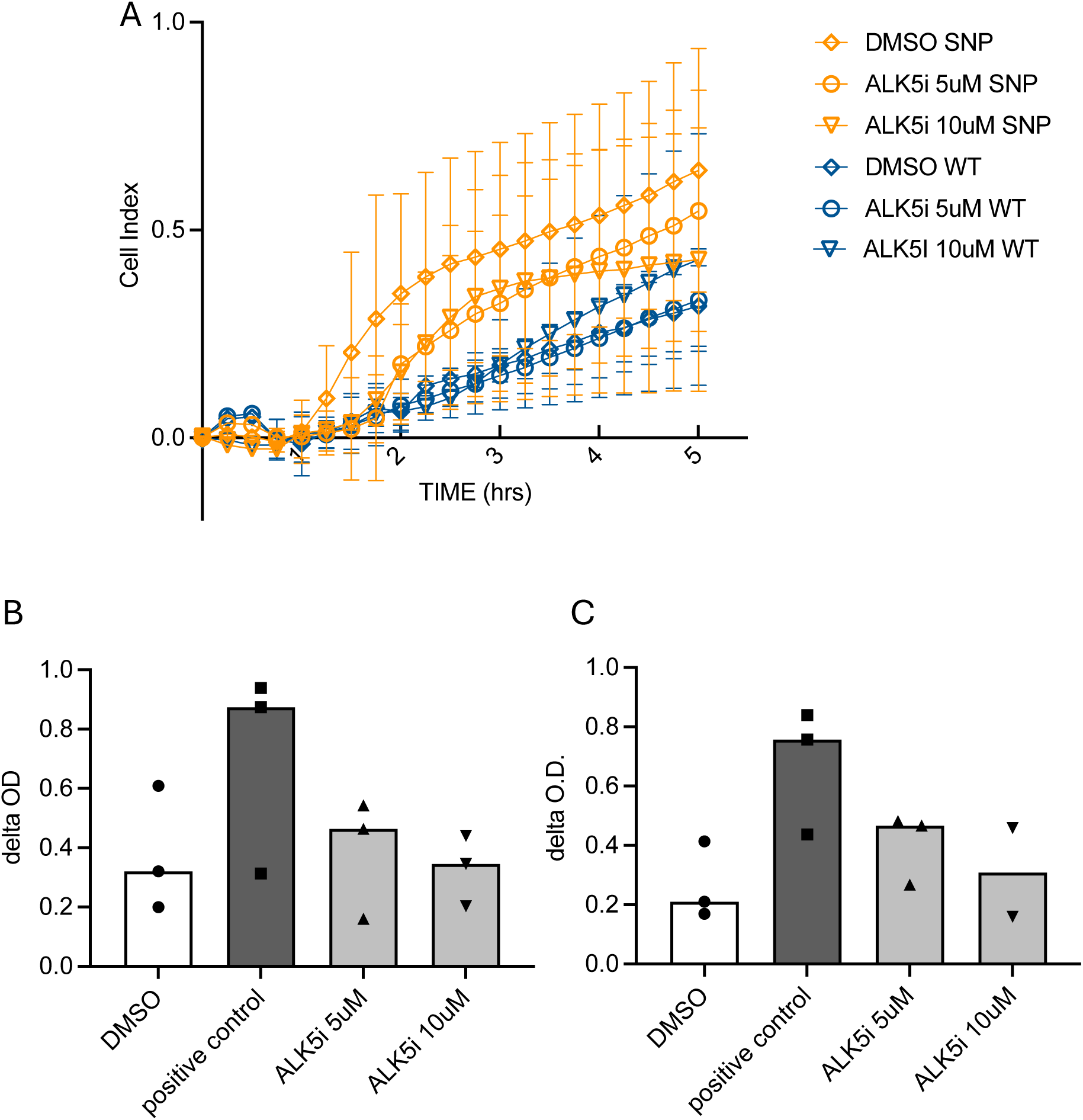
Effect of ALK5 inhibition on cell adhesion and cytotoxicity in iHBECs. (A) Real-time cell impedance measurements of rs62025270-bearing and WT iHBECs treated with an ALK5 inhibitor (5 or 10 µM) or DMSO control using the xCELLigence RTCA system. Data are presented as cell index over a 5-hour period (n = 3). (B) Lactate dehydrogenase (LDH) levels in rs62025270-bearing and WT iHBECs treated with an ALK5 inhibitor (5 or 10 µM) or DMSO control (n = 3).

## Notes

Conflict of Interest R.G.J. reports honoraria from Boehringer Ingelheim, Chiesi, Roche, PatientMPower, AstraZeneca, GSK, and consulting fees from AbbVie, AdALta, Apollo Therapeutics, Brainomix, Bristol Myers Squibb, Chiesi, Cohbar, GlaxoSmithKline Pliant, RedX. A.E.J is founder and shareholder of Alevin Therapeutics. I.D.S reports honoraria from patientMpower and is sole director and statistical consultancy of IndigoSigma Insights Ltd. L.O. is currently employed by Novartis Pharmaceuticals Australia. B.L., J.M., S.G., H.Y.Y., I.K., L.Z., E.P., G.C., R.C.C, A.K.R. declare no competing interests.

### Competing Interest Statement

R.G.J. reports honoraria from Boehringer Ingelheim, Chiesi, Roche, PatientMPower, AstraZeneca, GSK, and consulting fees from AbbVie, AdALta, Apollo Therapeutics, Brainomix, Bristol Myers Squibb, Chiesi, Cohbar, GlaxoSmithKline Pliant, RedX. A.E.J is founder and shareholder of Alevin Therapeutics. I.D.S reports honoraria from patientMpower and is sole director and statistical consultancy of IndigoSigma Insights Ltd. L.O. is currently employed by Novartis Pharmaceuticals Australia. B.L., J.M., S.G., H.Y.Y., I.K., L.Z., E.P., G.C., R.C.C, A.K.R. declare no competing interests.

